# Lectin-like Intestinal Defensin Inhibits 2019-nCoV Spike binding to ACE2

**DOI:** 10.1101/2020.03.29.013490

**Authors:** Cheng Wang, Shaobo Wang, Daixi Li, Xia Zhao, Songling Han, Tao Wang, Gaomei Zhao, Yin Chen, Fang Chen, Jianqi Zhao, Liting Wang, Wei Sun, Yi Huang, Yongping Su, Dongqing Wei, Jinghong Zhao, Junping Wang

## Abstract

The burgeoning epidemic caused by novel coronavirus 2019 (2019-nCoV) is currently a global concern. Angiotensin-converting enzyme-2 (ACE2) is a receptor of 2019-nCoV spike 1 protein (S1) and mediates viral entry into host cells. Despite the abundance of ACE2 in small intestine, few digestive symptoms are observed in patients infected by 2019-nCoV. Herein, we investigated the interactions between ACE2 and human defensins (HDs) specifically secreted by intestinal Paneth cells. The lectin-like HD5, rather than HD6, bound ACE2 with a high affinity of 39.3 nM and weakened the subsequent recruitment of 2019-nCoV S1. The cloak of HD5 on the ligand-binding domain of ACE2 was confirmed by molecular dynamic simulation. A remarkable dose-dependent preventive effect of HD5 on 2019-nCoV S1 binding to intestinal epithelial cells was further evidenced by *in vitro* experiments. Our findings unmasked the innate defense function of lectin-like intestinal defensin against 2019-nCoV, which may provide new insights into the prevention and treatment of 2019-nCoV infection.

## Introduction

Viral infection is still a major cause of human diseases despite the great improvements in medical care and public health. Recently, a novel pathogenic virus named novel coronavirus 2019 (2019-nCoV) or severe acute respiratory syndrome coronavirus 2 (SARS-CoV-2) emerged worldwide^1^, bringing about a pneumonia epidemic due to human-to-human transmission^2^. The clinical manifestations of 2019-nCoV-infected patients are similar to those of patients infected by severe acute respiratory syndrome coronavirus (SARS-CoV) and Middle East respiratory syndrome coronavirus (MERS-CoV) rife in 2003 and 2012, respectively^3,4^. Respiratory failure, septicopyemia, cardiac failure, hemorrhage, and kidney failure are the primary causes leading to the death of patients infected by 2019-nCoV^5^.

Coronavirus owns enveloped positive-stranded RNA encoding the spike (S) protein, envelope protein, membrane protein, and nucleoprotein. S protein is composed of 1160-1400 amino acids and presents as a trimer before membrane fusion^6^. Many S proteins can be divided into S1 and S2 subunits after degradation by protease^7^. The S1 subunit contains a receptor-binding domain (RBD) and devotes to the initial viral attachment via recognizing a variety of host cell receptors, including aminopeptidase N (APN), angiotensin-converting enzyme-2 (ACE2), dipeptidyl peptidase 4 (DPP4), carcinoembryonic antigen-related cell adhesion molecule 1 (CEACAM1), and sugar^8,9,10^. Since the sequence of 2019-nCoV S protein is 76.47% homology to that of SARS-CoV^11^, ACE2, which is identified as the crucial target for SARS-CoV, was also considered the host receptor of 2019-nCoV^12^. In fact, it was reported that host cells expressed ACE2, instead of APN and DPP4, were sensitive to 2019-nCoV^13,14^. *In vitro* binding assay demonstrated that ACE2 enabled a potent interaction with 2019-nCoV S protein with an affinity of 14.7 nM^15^, which is 10-to 20-fold higher than that of ACE2 binding to SARS-CoV S protein.

In human, ACE2 is remarkably abundant on lung alveolar epithelial cells and small intestine enterocytes^16,17^. As known, human intestinal epithelium encompasses approximately 200 m^2^ of surface area^18^. Therefore, human intestine is conceivably susceptible to 2019-nCoV. However, the occurrence of intestinal symptoms is lower than that of respiratory symptoms as a whole. In 38 patients infected by 2019-nCoV, only one diarrhea case was found^19^. Recent meta-analysis supported that digestive symptoms such as diarrhea, nausea, and vomiting were relatively rare compared with fever and cough^20^. These data indicate that intestinal epithelium is actually not so easy to be infected by 2019-nCoV as expected. The underlying reason is desirable to be explored.

It is well known that intestinal epithelium directly interacts with the external environment and encounters trillions of microorganisms including various viruses. To cope with the microbial threat, intestinal cells have evolved to produce a plenty of antimicrobial peptides (AMPs) to strengthen the mucosal barrier^21^. Defensin is a group of small (2-5 KD) and amphiphilic AMPs. Human defensins (HDs) are divided into two subgroups, α and β, based on the amino acid sequence homology and disulfide connectivity. Paneth cells, the major secretory cells located in the crypts of small intestine, specifically secrete two kinds of α defensins (HD5 and HD6)^22^. As reported, HD5 is the most abundant α defensin in intestine^23^. Due to a lectin-like ability, HD5 can effectively bind lipids and glycosylated proteins^24,25^. Because 2019-nCoV spike and ACE2 are both glycosylated proteins, there is a high possibility for HD5 to interact with 2019-nCoV spike or ACE2 or both of them and thereby disturb viral entry into host cells.

In the present study, we demonstrated that HD5 displayed a strong ability to prevent 2019-nCoV S1 to bind to intestinal epithelial cells by blocking the ligand-binding domain (LBD) of ACE2. Our findings unmasked the innate defense function of intestinal HD5 against 2019-nCoV, which may benefit the on-going development of preventive and therapeutic drugs.

## Results and Discussion

Intestinal defensins are concentrated in the mucus layer after being secreted by Paneth cells and can contact with ACE2 receptor that locates on the brush border of the enterocytes^16,21,26^. To gain an insight into the molecular interaction, we analyzed the recruitments of HD5 and HD6 to ACE2 by biolayer interferometry (BLI). The affinity of HD5 binding to recombinant human ACE2 immobilized on AR2G biosensors was 39.3 nM (Figure 1A), whereas no association signal was observed for HD6 (Figure 1B). The binding affinity of HD5 was much higher than that of a hexapeptide inhibitor, ^438^YKYRYL^443^, derived from the RBD of SARS-CoV spike protein (46 μM)^27^. We previously discovered that HD5 is reduced to linear peptide, HD5_RED_, under the catalysis of thioredoxin system *in vivo*^28,29^. Interestingly, BLI revealed that similar to HD6, HD5_RED_ failed to bind ACE2 (Figure 1C). The structure-dependent interaction of HD5 with ACE2 is consistent with that cysteine replacement impairs the bactericidal and immunoregulatory efficiencies of HD5^30,31^, which expands the known roles of disulfide pairings in keeping the conformation of defensins.

**Figure 1.**
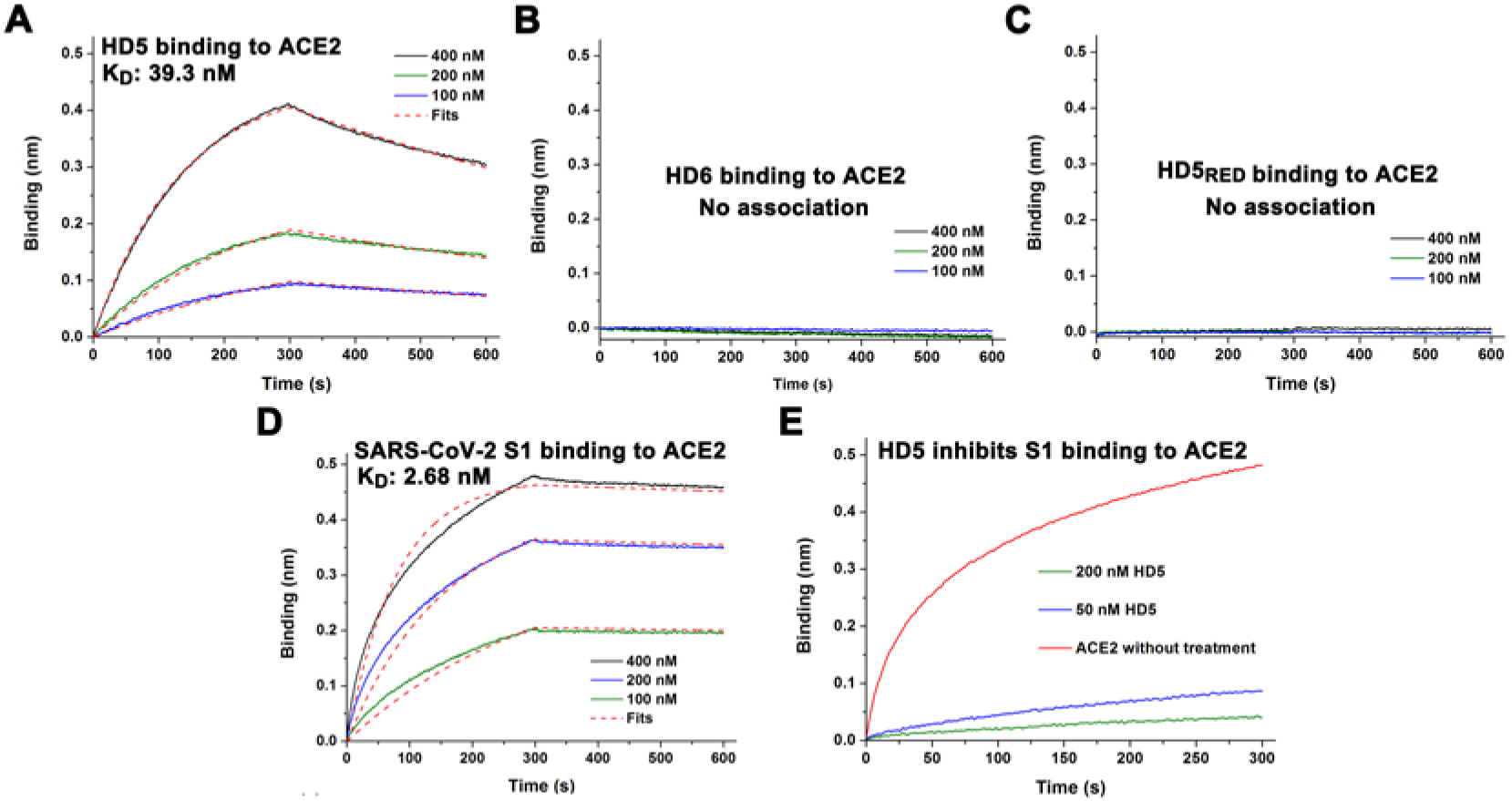
HD5 binds human ACE2 with a high affinity and inhibits 2019-nCoV S1 recruiting to ACE2. (A) Binding kinetics for HD5 and ACE2 loaded on AR2G biosensors. Fits of the data to a 1:1 binding model are shown with red dashes. Times for association and disassociation are both 300 s. (B) HD6 and HD5_RED_ (C) binding assay. (D) Binding kinetics for S1 and ACE2. (E) BLI-based ACE2 blocking experiment. The binding signals of 400 nM S1 to ACE2 coated with 200 and 50 nM HD5 are recorded for 300 s.

The interaction between 2019-nCoV S1 and ACE2 was measured as well. The purified S1 expressed by human HEK293 cells intensively bound ACE2 with an affinity of 2.68 nM (Figure 1D). This assay conforms to the fact that virus forwardly contact ACE2 located on the cell surface and may avoid the steric hindrance of a receptor binding to its ligand. We reversely loaded S1 and measured the recruitment of ACE2. As shown in Figure S1, the binding thickness was low, whereas the affinity (3.8 nM) was comparable to that of S1 binding to ACE2. Also, we noticed that the S1-ACE2 affinity was higher than those of S-ACE2 and RBD-ACE2 interactions^15,32^. The discrepancy may be attributed to different proteins employed in different experiments.

HD5 is constitutively expressed *in vivo* and acts as a peptide “ranger” regulating the intestinal microbiota^33^. The storage volume of HD5 in Paneth cells has been estimated 90-450 mg/cm^2^ of ileal surface area^22^. In ileal fluid, HD5 content is quantified to 6-30 μg/mL^34^. Such a large dose of HD5 in intestine provides a possibility that it may capture ACE2 before 2019-nCoV lands to enterocytes, although the affinity of S1-ACE2 interaction is higher than that of HD5 binding to ACE2. To evaluate the effect of HD5 coating on ACE2 interacting with S1, a blocking experiment was conducted by monitoring the bindings of S1 to ACE2 covered with different doses of HD5. As shown in Figure 1E, HD5 lowered the thickness of S1 binding to ACE2, indicative of a protective role of this peptide on viral adhesion. As HD5 is lectin-like peptide with multivalent binding ability^24^, we asked if HD5 could bind and block S1. The affinity of HD5 binding to S1 was 82.8 nM (Figure 2A), as revealed by BLI. Nevertheless, HD5 had less of an effect on ACE2-S1 interaction (Figure 2B), possibly due to a binding site distant from the RBD. Recent studies have discovered some antibodies and peptides with high affinities binding to the spike^32,35^. These candidates are placed hopes on suppressing 2019-nCoV. Of note, our findings imply that the binding affinity alone is insufficient to predict the therapeutic potential of drugs.

**Figure 2.**
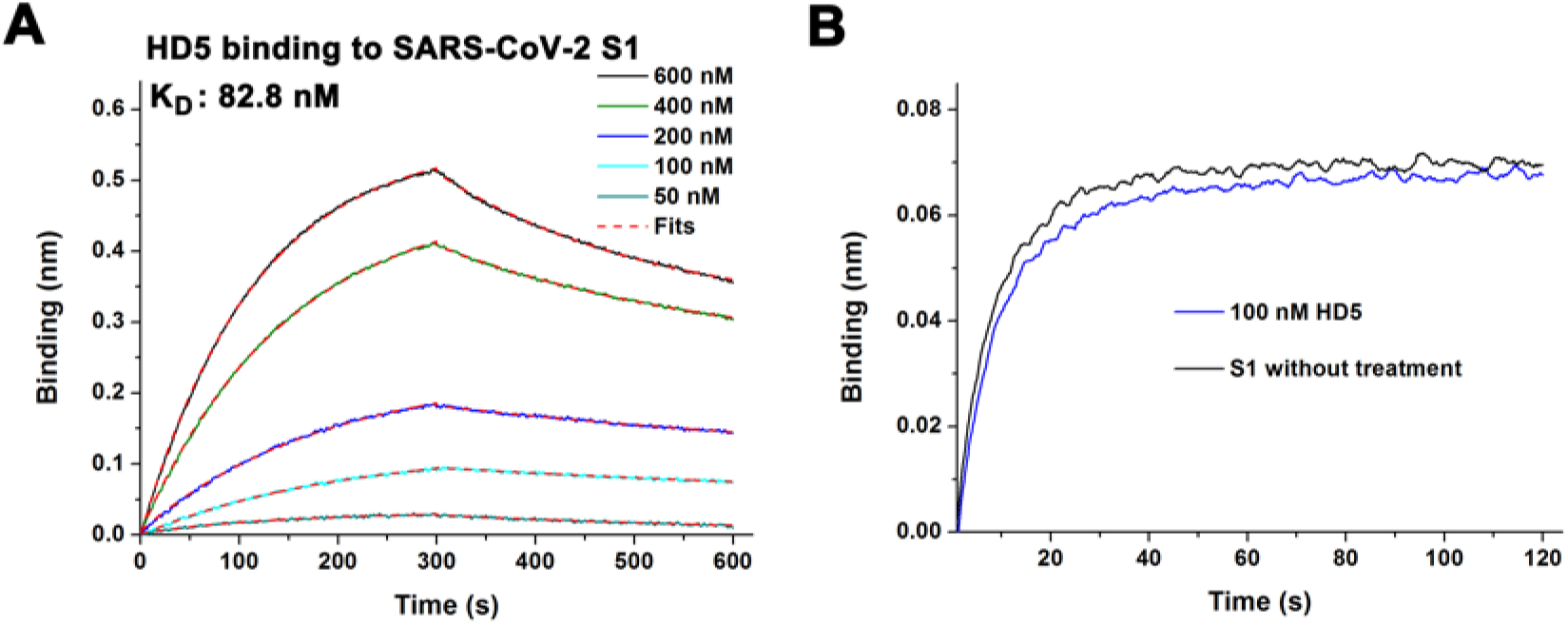
HD5 binds 2019-nCoV S1 but fails to inhibit S1 interacting with ACE2. (A) Binding kinetics for HD5 and S1 loaded on HIS1K biosensors. Fits of the data to a 1:1 binding model are shown with red dashes. (B) BLI-based S1 blocking assay. The binding signals of 100 nM ACE2 to S1 coated with 100 nM HD5 are recorded for 120 s.

X-ray crystal diffraction has resolved the detailed structure of 2019-nCoV RBD bound with ACE2 at 2.45 Å resolution, in which 20 residues of N-terminal ACE2 constitute the LBD^36^. For deeper insights into the ACE2 blocking of HD5, we carried out a molecular dynamic simulation, in which HD5 was docked onto the LBD of ACE2. After 20 ns of simulation, the complex conformation kept stable with residue fluctuations smaller than 0.11 nm in five trajectories relative to the starting coordinates (Figure S2A). HD5 formed many potent H-bonds with ACE2, which average number is 5-6 and the max number is 12 in an average distance of 2.825 Å and an average angle of 14.5° during 20-ns simulation (Figure S2B-D). Additionally, three electrostatic interactions (HD5-Arg^6^ ~ ACE2-Glu^23^, HD5-Glu^21^ ~ ACE2-Lys^31^, HD5-Glu^21^ ~ ACE2-His^34^) and three hydrophobic interactions (HD5-Ala^1^ ~ ACE2-Leu^79^, HD5-Tyr^4^ ~ ACE2-Phe^28^, HD5-Tyr^4^ ~ ACE2-Leu^79^) took part in the interface stabilization (Figure 3A), resulting in a net free binding energy of −1866.53 kJ/mol.

**Figure 3.**
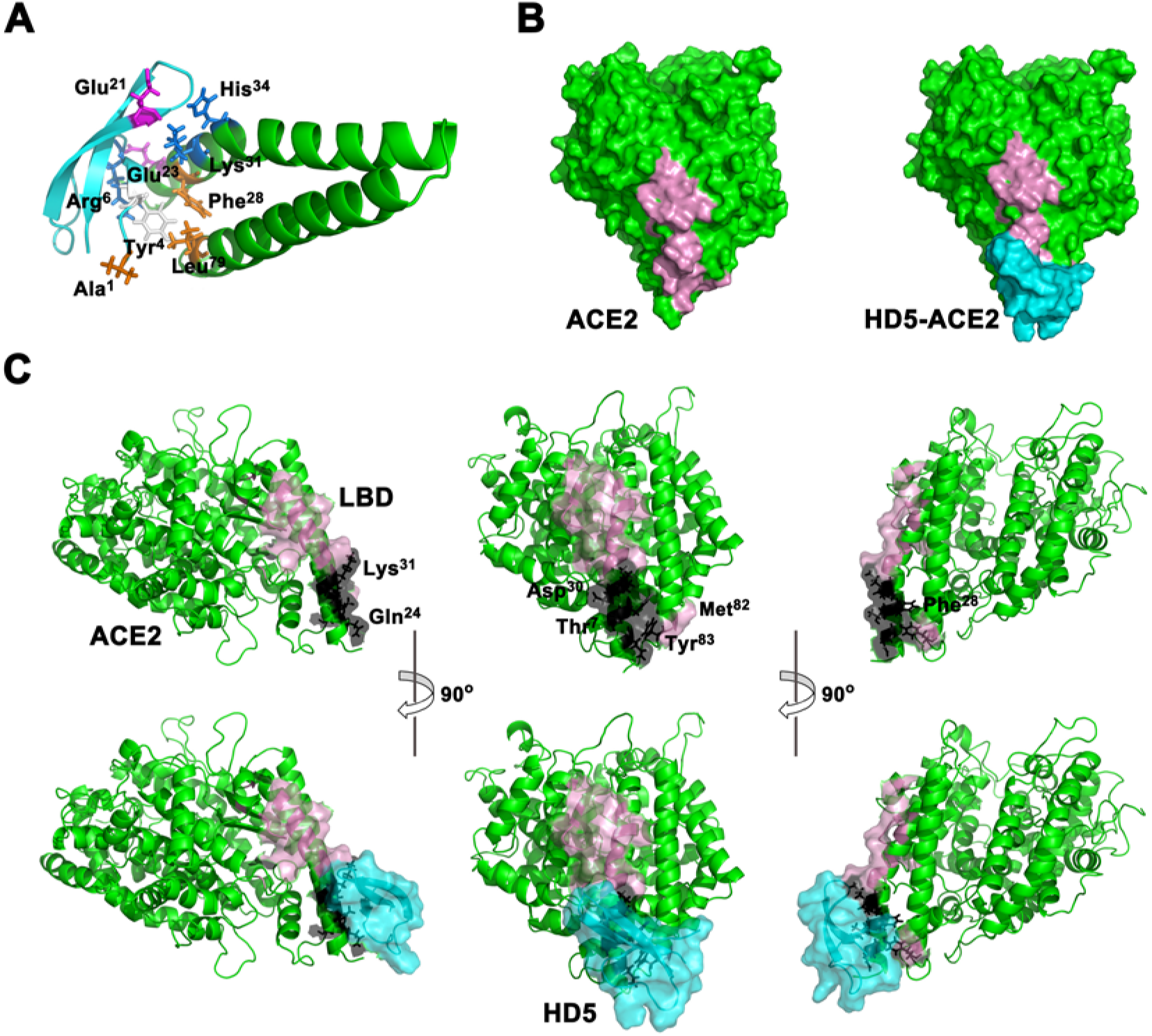
Structural analysis for ACE2 bound with HD5 after molecular dynamic simulation. (A) Ribbon diagram of the binding interface. HD5 colored cyan is composed of 32 residues constrained by three disulfide bonds, displaying as a three-stranded antiparallel β-sheet conformation in steric. ACE2 α-helix 1 and 2 (Ser^19^ to Leu^85^) are colored green. Residues contributing to the interface stabilization are presented as sticks. Positively and negatively charged residues are highlighted in blue and magenta, respectively. Hydrophobic residues are orange. The uncharged hydrophilic residue is shown in gray. (B) Topological view of ACE2 bound with HD5. ACE2 and its LBD are colored green and pink, respectively. (C) Stereoview of the shielding effect of HD5 on ACE2 LBD. Residues in the LBD cloaked by HD5 are colored black.

Driven by the potent intermolecular interaction, HD5 tightly attached the surface of ACE2 LBD (Figure 3B) and cloaked Thr^7^, Gln^24^, Asp^30^, and Lys^31^ on α-helix 1 and Tyr^83^ on loop 2 (Figure 3C). The exposure of Met^82^ was partially shielded. Among the blocking sites, Asp^30^, Lys^31^, and Met^82^ might be more important than the others, as mutations of Asp^30^ and Lys^31^ caused a marked loss of interaction between ACE2 and SARS-CoV S1^37^, and Met^82^ mutations in rat and mouse lowered the entry of SARS-CoV^38^. Considering the sequence difference between SARS-CoV and 2019-nCoV^11^, further research is required for investigating whether these sites are critical determinants for 2019-nCoV binding to ACE2.

To validate the inhibitive effect of HD5 on 2019-nCoV S1 binding to ACE2, we further conducted a cell experiment, in which human intestinal epithelium Caco-2 cells were exposed to 8 μg/mL of 2019-nCoV S1. Confocal microscopy observed that S1 largely adhered to the cell surface in the absence of HD5 (Figure 4, Figure S3). Nevertheless, when cells were preincubated with 100 μg/mL of HD5, the recruitment of S1 was dramatically reduced. Western blot supported that HD5 markedly protected Caco-2 from the adherence of S1 (Figure 5A). The content of S1 binding to the cells preincuabted with HD5 was 3.4-fold lower than that binding to the cells without treatment (Figure 5B). Notably, the protection of HD5 was in a dose-dependent manner (Figure 5C). As shown in Figure 5D, HD5 could significantly decrease S1 binding to epithelial cells at concentrations as low as 10 μg/mL, which is within the range of HD5 concentration in intestine^22^. The protective effect of HD5 was also observed for human renal proximal tubular epithelial cells abundant in ACE2 (Figure S4)^16,17^. However, S1 pretreated with HD5 was still efficient to contact Caco-2, as revealed by either immunoblotting or immunofluorescence. These data are in line with the results shown in Figure 1E and 2B, demonstrating that HD5 inhibits 2019-nCoV S1 adhering to host cells possibly by blocking ACE2 but not S1.

**Figure 4.**
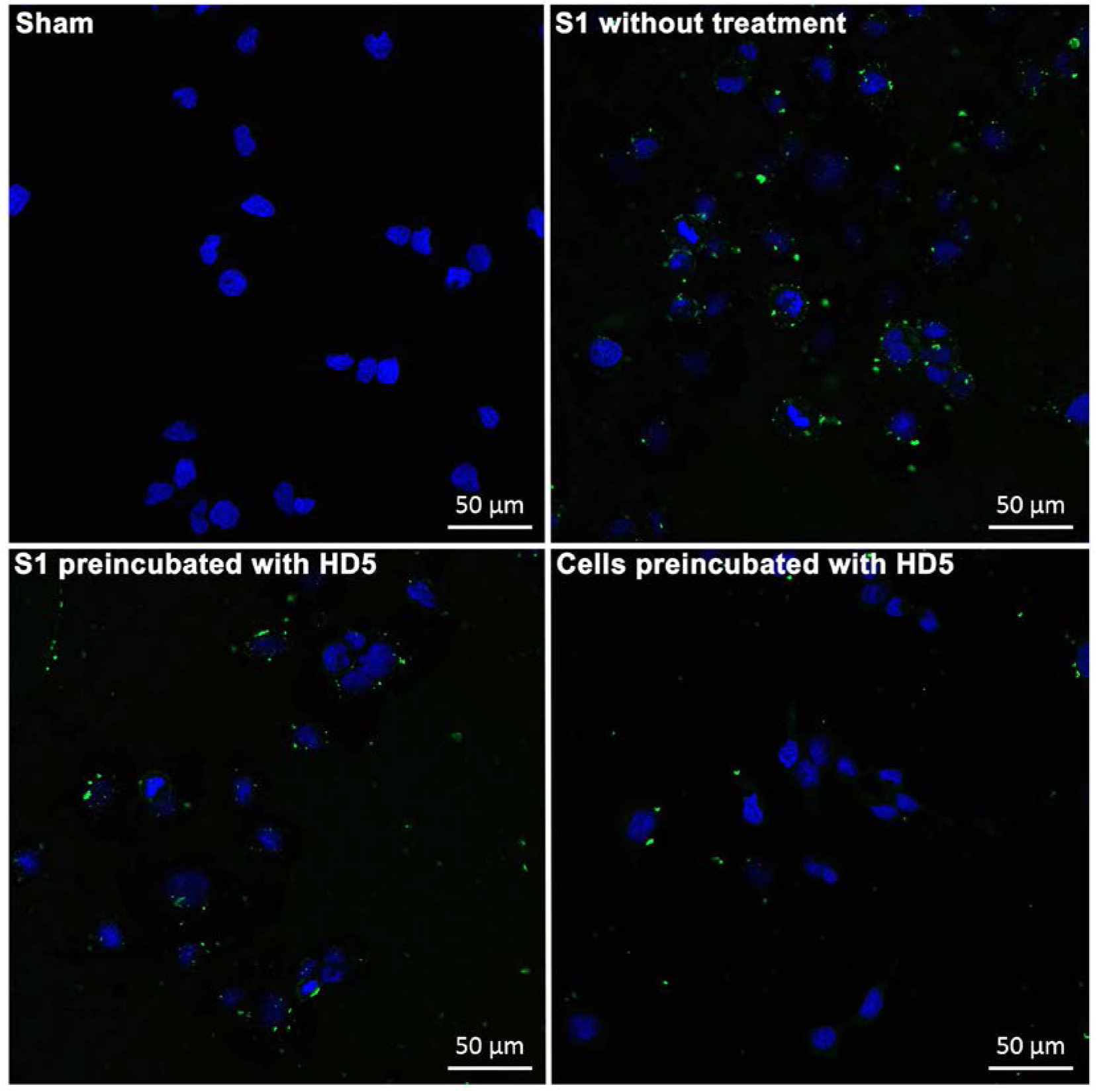
Immunofluorescence microscopy revealing the protection of HD5 on intestinal cells exposed to 2019-nCoV S1. S1 adhering to the cell surface is probed by a goat anti-rabbit Alexa Fluor 488 antibody (Green). Nuclei are stained using DAPI (blue). Scale bar indicates 50 μm.

**Figure 5.**
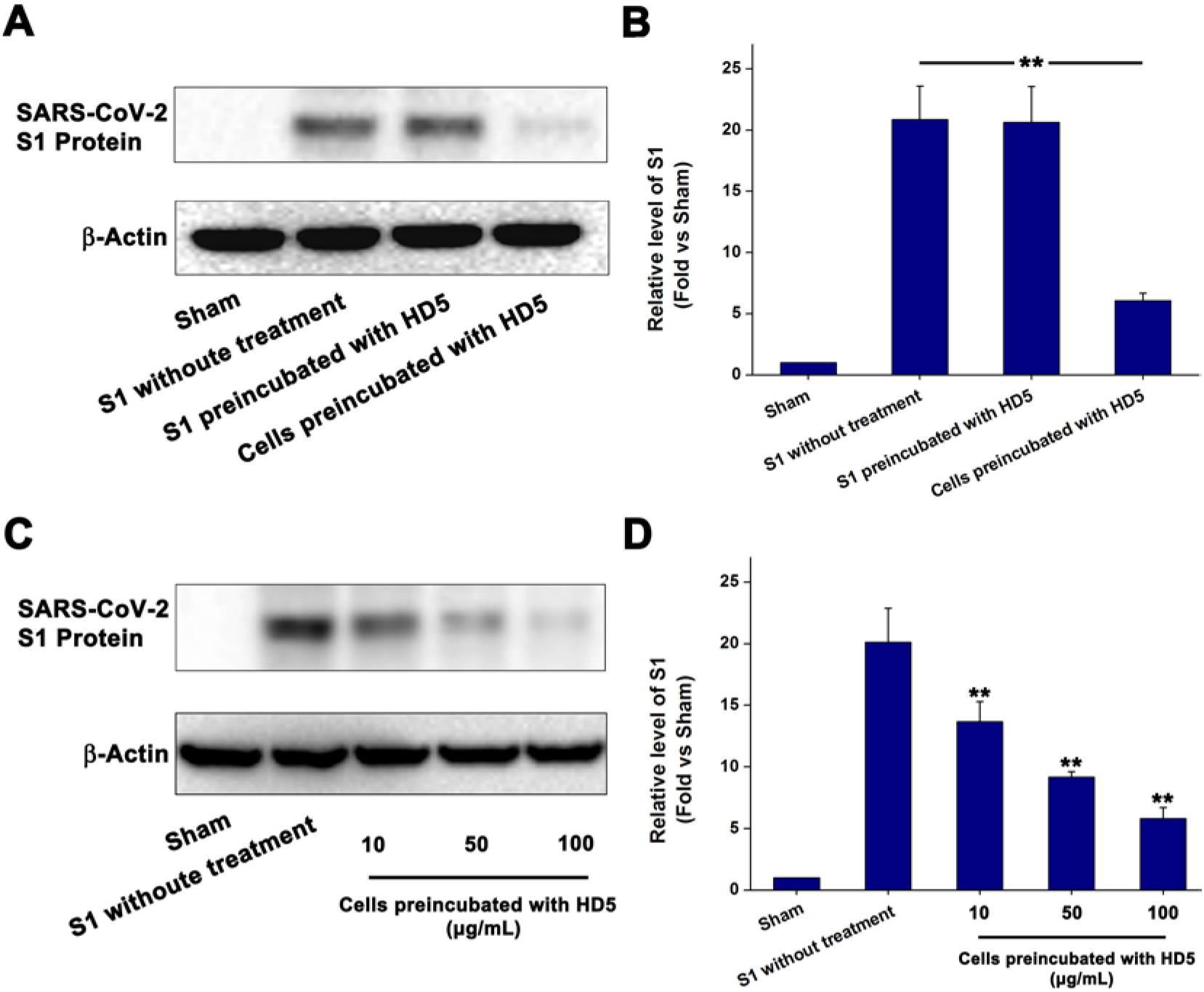
Western blot demonstrating the inhibition of HD5 on 2019-nCoV S1 binding to intestinal cells. (A) Caco-2 cells preincubated with 100 μg/mL of HD5 are less efficient to recruit S1 than those without treatment. A total of 25 μg of each protein sample is resolved by 10% SDS-PAGE. (B) Histogram presenting S1 levels obtained by gray scanning relative to the reference. **, *P* < 0.05. (C) Shown is the protein band of S1 binding to Caco-2 treated with increasing concentrations of HD5. (D) HD5 protects intestinal cells from the adherence of S1 in a dose dependent manner. **, *P* < 0.05, relative to the group of S1 without treatment.

Collectively, we found for the first time that lectin-like intestinal defensin can protect host cells against 2019-nCoV entry. Paneth cell-secreted HD5 efficiently bound and blocked ACE2 which locates on the surface of intestinal epithelial cells, lowering the recruitment of 2019-nCoV S1 (Figure 6). It is an interesting explanation to the clinical phenomenon that few intestinal symptoms are observed in patients infected by 2019-nCoV^19^. Due to the immature Paneth cells and deficiency in HD5^39^, it is might also be a potential reason for the infection of a newborn 30 h after birth^40^. Similarly, adult patients receiving small intestine transplantation or suffering from inflammatory bowel diseases such as the Crohn’s disease, whose HD5 content are deficient *in vivo*^41,42^, might be more susceptible to 2019-nCoV than the healthy. More importantly, as there is a shortage of effective drugs to prevent and treat 2019-nCoV infection, we think that it may be a useful strategy to increase the content of HD5 *in vivo* by immunoregulation or exogenous supplement of HD5 as we recently described^43^.

**Figure 6.**
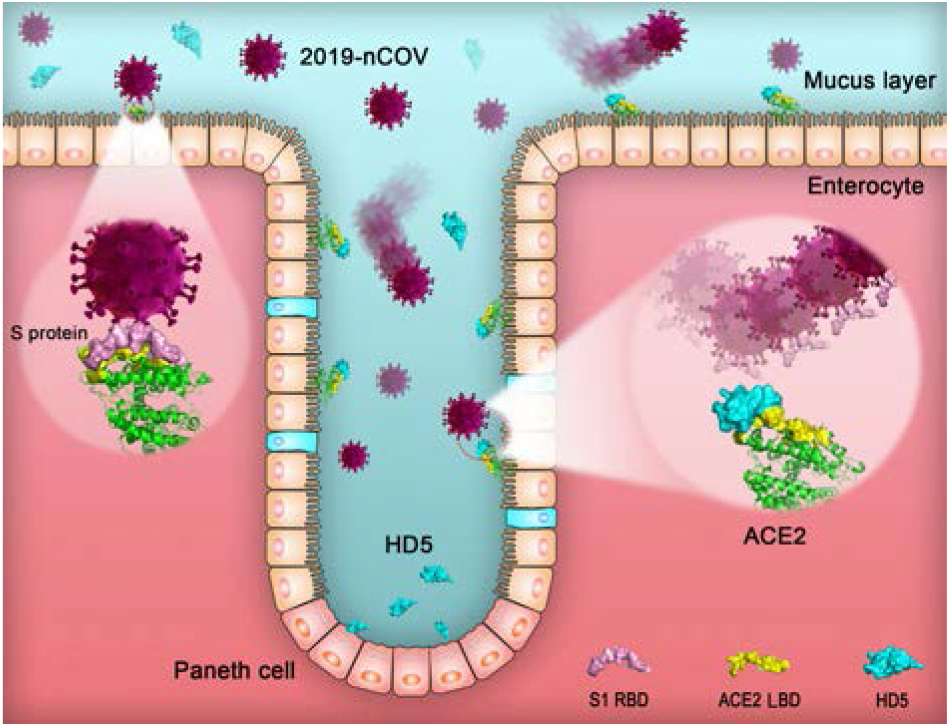
Schematic illustration of the Paneth cell-mediated host defense against 2019-nCoV. Human Paneth cells specifically secrete a lectin-like peptide named HD5, the most abundant α defensin in intestine. HD5 intensively binds and blocks epithelial ACE2, which is located on the brush border of enterocytes and is a recognized receptor for 2019-nCoV spike, thus weakening the viral adhesion and exerting a protective effect.

## Methods

### Peptide synthesis

Peptides were prepared by Chinese Peptide Company (Hangzhou, Zhejiang Province, CHN) and purified with an Agilent (Beijing, CHN) 1260 HPLC system equipped with a Phenomenex/Luna C18 column (5 μm, 4.6×150 mm). The purities and molecule masses of peptides determined by reverse-phase high performance liquid chromatography (RP-HPLC) and electrospray ionization mass spectrometry, respectively, were shown in Table S1.

### Biolayer interferometry (BLI)

The bindings of peptides and 2019-nCoV S1 (40591-V08H, Sino Biological) to recombinant human ACE2 (10108-H02H, Sino Biological, Beijing, CHN) was measured using Forte Bio’s “Octet Red 96” BLI (Pall Life Sciences, Port Washington, NYC, US). ACE2 (15 μg/mL) was immobilized on AR2G biosensors activated by 1-ethyl-3-[3-dimethylaminopropy] carbodiimide hydrochloride (EDC, E1769, Sigma, Shanghai, CHN) and sulfo-N-hydroxysulfosuccinimide (s-NHS, 56485, Sigma, Shanghai, CHN). Peptides and S1 was prepared in PBS with concentrations of 400, 200, and 100 nM. To determine the recruitment of HD5 and ACE2 to S1, S1 was loaded on HIS1K biosensors at 15 μg/mL. HD5 was prepared in PBS with concentrations of 600, 400, 200, 100, and 50 nM. ACE2 was diluted to 75, 50, and 40 nM in PBS. Association and disassociation, 300 s for each, were performed at a shaking speed of 1000 rpm. The results were processed using Fortebio Data Analysis 7.0 software. The equilibrium dissociation constant (K_D_) was generated by a 1:1 fitting model.

BLI-based blocking assay was conducted by monitoring the binding thickness of S1-ACE2 interaction with the interference of HD5. S1 and ACE2 were loaded on HIS1K and AR2G biosensors, respectively, at a concentration of 15 μg/mL. The immobilized ACE2 was incubated with 200 and 50 nM HD5 for 300 s at 30°C. After a 300 s of disassociation, the signals of 400 nM S1 binding to ACE2 were recorded for another 300 s. Otherwise, the immobilized S1 was covered with 100 nM HD5 for 300 s. The bindings of 100 nM ACE2 to S1 were recorded for 300 s after peptide disassociation.

### Molecular dynamic simulation

Molecular dock between HD5 (PDB: 1ZMP^44^) and the LBD of ACE2 (PDB: 6ACG^45^) was performed on ZDOCK server^46^. The Gromacs 2020 software package^47^, AMBER99SB-ILDN force field, and TIP3P water model were applied for the simulation with a time step of 2 fs. Firstly, 1000-step minimization was carried out. Then, for fully relaxation, four 1-ns pre-equilibration simulation with restrained coordinates of the atoms belonging to the heavy atoms, main chain, backbone, and C-α, respectively, were performed step by step. Finally, each of five production simulation with isothermal-isobaric (NPT) ensemble at 1 atm and 298 K was performed for 20 ns. The binding free energy was calculated using the molecular mechanics-Poisson-Boltzmann surface area (MM-PBSA) method with g_mmpbsa procedure for Gromacs^48, 49^. The binding free energy was calculated using 500 snapshots sampled every 10 ps from total 5 ns trajectory.

### Immunofluorescence microscopy

Caco-2 and HK-2 obtained from the cell bank of Chinese Academy of Sciences (CAS, Shanghai) and cultured in Dulbecco’s modified Eagle medium (DMEM, Gibco, Thermo Fisher Scientific, Shanghai, CHN) containing 10% foetal bovine serum (FBS, Gibco) were seeded into a 12-well plate with sterile glass slides at a density of 2×10^5^ cells/well. Cells cultured overnight were washed with PBS and preincubated with 100 μg/mL of HD5 at 37 °C for 15 min, followed by an addition of 8 μg/mL of S1. Co-incubation was further performed at 4 °C for 1 h. After three times of wash with PBS, cells were fixed in 4% paraformaldehyde. A primary anti-spike rabbit monoclonal antibody (40150-R007, Sino Biological, 1:100) and a goat anti-rabbit secondary antibody (Alexa Fluor 488, Invitrogen, Thermo Fisher Scientific) were employed to stain 2019-nCoV S1. Nuclei were stained with 2-(4-amidinophenyl)-6-indolecarbamidine dihydrochloride (DAPI, C1002, Beyotime, Shanghai, CHN). A Zeiss LSM 780 NLO confocal microscope was employed to observe the cells.

### Western blot

Caco-2 cells were seeded into a 6-well plate at a density of 1×10^6^ cells/well. Cells preincubated with different concentrations of HD5 (10, 50, and 100 μg/mL) at 37 °C for 15 min were exposed to 20 μg/mL of 2019-nCoV S1 containing His-tag at 4 °C for 1 h. After three times of wash with PBS, cells were collected and processed with RIPA lysis and extraction buffer (89900, Thermo Fisher Scientific). S1 content was measured by determining the band thickness as we previously described^50^. A primary anti-His-tag mouse monoclonal antibody (AF5060, Beyotime, 1:1000) and a goat anti-mouse secondary antibody (A0216, Beyotime, 1:2000) were employed to detect S1. β-actin determined by a mouse monoclonal antibody (AA128, Beyotime, 1:1000) was used as a reference.

## Supporting Information

Figures and Tables are as noted in text. This material is available via the internet at https://www.biorxiv.org/.

## Acknowledgement

This work was supported by grants from the National Natural Science Foundation of China (Nos. 81725019 and 81703395), the program for scientific and technological innovation leader of Chongqing, the Natural Science Fund of Chongqing City (cstc2019jcyj-msxmX0011), and the frontier specific projects of Xinqiao Hospital (2018YQYLY004). We have no competing financial interests to declare.

