## Supplementary material for "Lectin-like Intestinal Defensin Inhibits 2019-nCoV Spike binding to ACE2": Table S1, Figure S1-4

**Contents**

Table S1. Sequence, purity, and mass of the peptides

Figure S1. Binding kinetics for ACE2 and S1 loaded on HIS1K biosensors

Figure S2. Molecular dynamic simulation analyzing HD5 interacting with ACE2

Figure S3. Immunofluorescence microscopy revealing the effect of HD5 on Caco-2 cells

Figure S4. Immunofluorescence microscopy revealing the protection of HD5 on HK-2 cells exposed to 2019-nCoV S1

**Table S1. Sequence, purity, and mass of the peptides.**

| Peptide | Sequence | Purity [a] | Theoretical mass [b] | Measured mass [c] |
|---|---|---|---|---|
| HD5 | ATCYCRTGRCATRESLSGVCEISGRLYRLCCR | 96.0% | 3582.2 | 3582.3 |
| HD6 | AFTCHCRRSCYSTEYSYGTCTVMGINHRFCCL | 95.7% | 3708.3 | 3707.6 |
| $HD5_{RED}$ | Linear-ATCYCRTGRCATRESLSGVCEISGRLYRLCCR | 96.2% | 3588.2 | 3588.2 |

[a] Measured by reverse-phase high performance liquid chromatography. The chromatographic data were obtained at 40˚C on a Phenomenex/Luna C18 (2) column (5 µm, 4.6×150 mm) applying a linear gradient of 20-40% buffer B (buffer A: 0.1% trifluoroacetic acid in water; buffer B: 0.09% trifluoroacetic acid in (80% acetonitrile plus 20% water)) at a flow rate of 1 mL $min^{-1}$ over 20 min.

[b] Calculated on http://web.expasy.org/protparam/.

[c] Measured by electrospray ionization mass spectrometry (Chiron Mimotopes, Victoria, Australia).

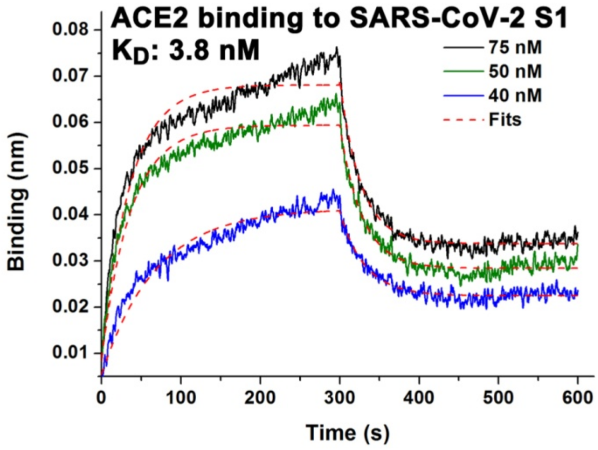


**Figure S1. Binding kinetics for ACE2 and S1 loaded on HIS1K biosensors.** Fits of the data to a 1:1 binding model are shown with red dashes. Times for association and disassociation are both 300 s.

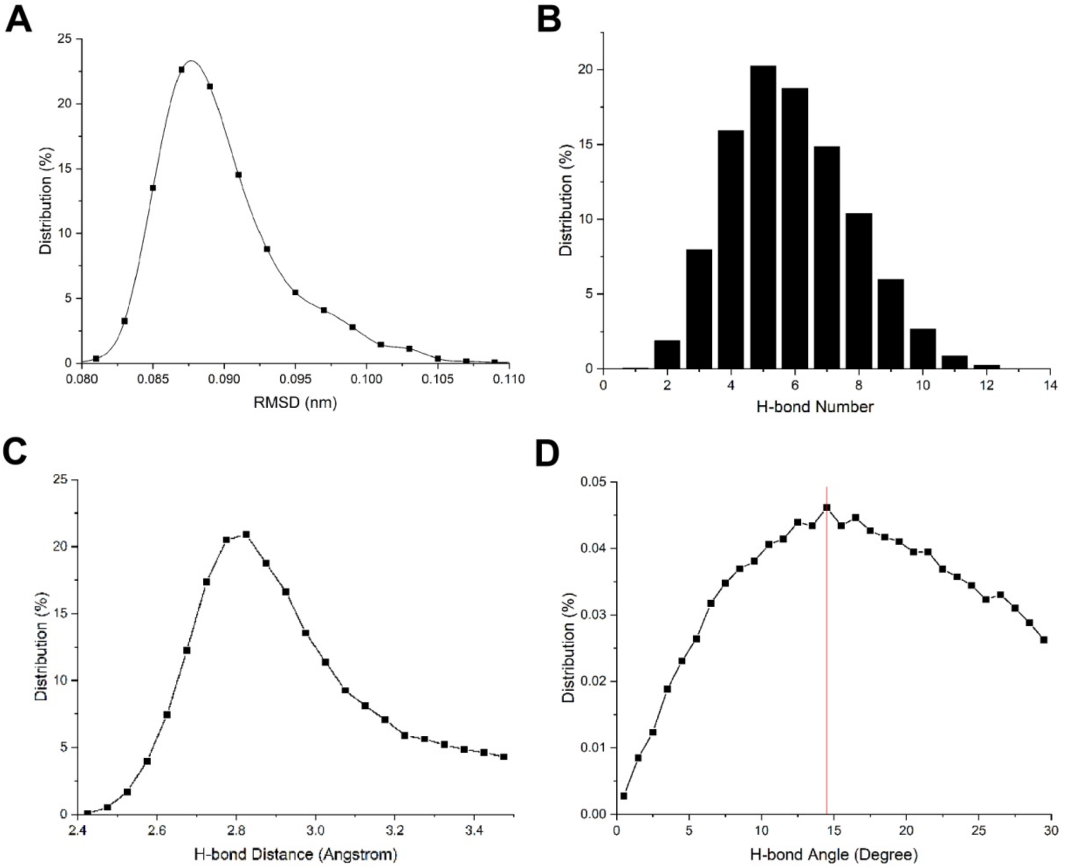


**Figure S2. Molecular dynamic simulation analyzing HD5 interacting with ACE2.** (A) Root-mean-square deviation (RMSD) of residue fluctuations. (B) Number distribution of H-bonds. The average number is 5-6. The most number reaches 12. (C) Distance distribution of H-bonds. The average distance is 2.825 Å. (D) Angle distribution of H-bonds. The average Angle is 14.5$^{o}$ .

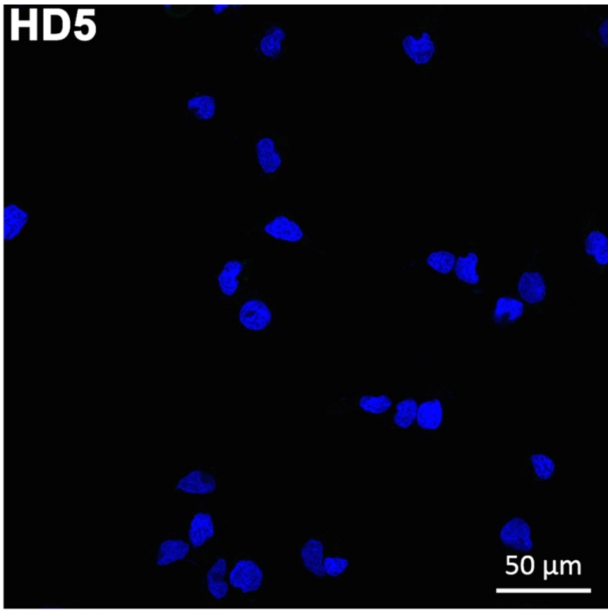


**Figure S3. Immunofluorescence microscopy revealing the effect of HD5 on Caco-2 cells.** Cells are incubated with 100 µg/mL of HD5 in PBS at 4 $^{o}$C for 1 h. Nuclei are stained using DAPI (blue). Scale bar indicates 50 µm.

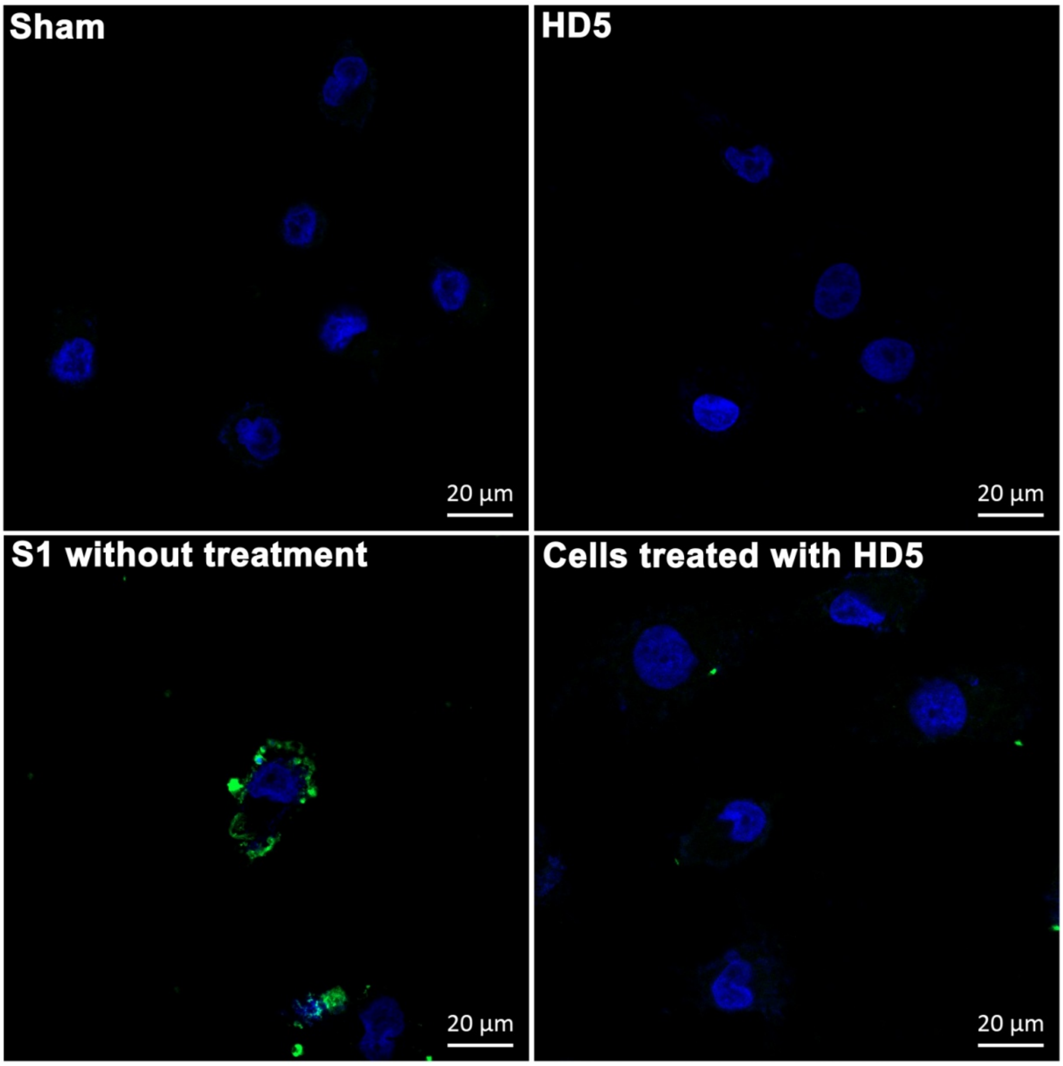


**Figure S4. Immunofluorescence microscopy revealing the protection of HD5 on HK-2 cells exposed to 2019-nCoV S1.** S1 adhering to the cell surface is probed by a goat anti-rabbit Alexa Fluor 488 antibody. Nuclei are stained using DAPI (blue). Scale bar indicates 20 μm.
